# Microtubule search-and-capture model evaluates the effect of chromosomal volume conservation on spindle assembly during mitosis

**DOI:** 10.1101/2023.04.08.536118

**Authors:** Pinaki Nayak, Saptarshi Chatterjee, Raja Paul

**Affiliations:** School of Mathematical and Computational Sciences, Indian Association for the Cultivation of Science, Kolkata-700032, India; Wallace H. Coulter Department of Biomedical Engineering, Georgia Institute of Technology and Emory University, Atlanta, Georgia 30332, USA

**Author notes:** contributed equally to this paper.

## Abstract

Variation in the chromosome numbers can arise from the erroneous mitosis or fusion and fission of chromosomes. While the mitotic errors lead to an increase or decrease in the overall chromosomal substance in the daughter cells, fission and fusion keep this conserved. Variations in chromosome numbers are assumed to be a crucial driver of speciation. For example, the members of the muntjac species are known to have very different karyotypes with the chromosome numbers varying from 2*n* = 70 + 3*B* in the brown brocket deer to 2*n* = 46 in the Chinese muntjac and 2*n* = 6*/*7 in the Indian muntjac. The chromosomal content in the nucleus of these closely related mammals is roughly the same and various chromosome fusion and fission pathways have been suggested as the evolution process of these karyotypes. Similar trends can also be found in lepidoptera and yeast species which show a wide variation of chromosome numbers. The effect of chromosome number variation on the spindle assembly time and accuracy is still not properly addressed. We computationally investigate the effect of conservation of the total chromosomal substance on the spindle assembly during prometaphase. Our results suggest that chromosomal fusion pathways aid the microtubule-driven Search and Capture of the kinetochore in cells with monocentric chromosomes. We further report a comparative analysis of the site and percentage of amphitelic captures, dependence on cell shape, position of the kinetochore in respect of chromosomal volume partitioning.

## I. INTRODUCTION

The process of cell division is one of the most complex and widely studied cell biological phenomena. More often than not, the mitotic spindle, primarily a microtubule-based self-assembling cellular machine, facilitates the process of cell division where the genetic material encoded in the chromosomes is segregated and passed on to the progeny cells from the mother [1, 2]. In general, during centrosomal spindle assembly, microtubule minus ends are anchored at two centrosomes serving as two spindle poles. During prometaphase, highly dynamic microtubule plus-ends progressively establish physical linkage with kinetochores-a complex multi-protein structure associated with the centromere region of the chromosomes, through a proposed search-and-capture process.

The concept of search-and-capture quite naturally entails a ‘searcher’ combing through the space for a ‘target’ to be captured. The microtubule search-and-capture hypothesis is fundamentally based on the microtubule dynamic instability [3–8]. According to this, microtubules growing in random directions repeatedly toggle between growth and shrinkage phases. The stochastic transition from growth to shrinkage is known as a catastrophe whereas the reverse transition is known as a rescue. Microtubule nucleation in random directions together with dynamic instability inherently enable the microtubules to search for ‘targets’ within the cellular geometry [4, 9, 10]. In the proposed microtubule search-and-capture model, when a microtubule plus end encounters a kinetochore, a microtubule-kinetochore connection is established-in other words, that particular kinetochore is captured. This model, in particular, elucidated two key aspects of mitotic spindle assembly-(a) microtubule-mediated capture of chromosomes and (b) the nature of microtubule-kinetochore attachments. The model, to a large extent, explained the rapid establishment of physical connections between all the chromosomes and spindle poles (chromosome capture) within minutes as assessed via experiments [9]. In this article, we termed the time elapsed in capturing all the kinetochores via centrosomal microtubules as the total capture time. However, mere capture of the chromosomes is a necessary but not sufficient condition for the progression of spindle assembly- the nature and accuracy of the microtubule-kinetochore attachments determine the fate of faithful, error-free chromosome segregation [11]. Since microtubules from both the spindle poles can attach to kinetochores, multiple types of attachments are possible [12]. Interestingly, barring one, all the other types are incorrect kinetochore-microtubule attachments which give rise to segregation defects. The biologically correct attachment is termed amphitelic attachment where each sister kinetochore is attached to microtubules extending solely from one of the two opposite spindle poles. On the other hand, the incorrect attachments are (a) syntelic attachments where both sister kinetochores are linked to the same spindle pole and there exists no connection between the other spindle pole and the sister kinetochores; (b) monotelic attachments where only a single kinetochore is connected to just one spindle pole and the other kinetochore remains unattached from both the spindle poles and (c) merotelic attachments where at least one kinetochore is attached to microtubules emanating from both the spindle poles. By definition, the formation of merotelic attachments requires more than one microtubule binding site on a kinetochore. Due to the complexity of the spindle assembly process, computational modeling is proven to be a useful tool to complement/support the experimental research focusing on various aspects of spindle dynamics across organisms [13–27]. To understand the speed and accuracy of the spindle assembly process, several variants of search-and-capture models incorporating microtubules and kinetochores in two-dimensional (2D) and three-dimensional (3D) geometries have been utilized to quantitatively identify the conditions and mechanisms that facilitate error-free segregation [9, 10, 28–36].

Our aim in this study is to examine the relationship between chromosome number and the partitioning of the total chromosomal content in the light of search-and-capture. Chromosome number varies widely across different species. For example, *Drosophila melanogaster* (fruit fly) has only 4 pairs of chromosomes, whereas *Caenorhabditis elegans* (worm), *Saccharomyces cerevisiae* (yeast), *Homo sapiens* (humans), and *Canis familiaris* (dog) have 6, 16, 23, and 39, respectively. Of note, the females of the Indian muntjac (*Muntiacus muntjak*), a placental mammal, commonly known as barking deer, contains the lowest known diploid chromosome number (2N = 6) with large kinetochores among mammals [37–40]. Muntjac species are thought to have descended from a common ancestor but their karyotypes are vastly different in each species. Intriguingly, while the Indian muntjac has a diploid number of 6 in the female and 7 in the male, the Chinese muntjac has a diploid number of 46 and the brown brocket deer has a diploid number of 70 + 3B. Chromosome painting experiments have suggested that the chromosomal substance of the 46 chromosomes of the Chinese muntjac is condensed into the 6/7 chromosomes of the Indian muntjac [41]. It is likely that the members of the muntjac species have evolved from their common ancestor through three plausible mechanisms: (a) chromosome fission, (b) chromosome fusion and, (c) redistribution of the chromosomal substance. Experimentally, reducing chromosome numbers through chromosome fusion has been achieved by karyotype engineering in yeast cells leading to reproductive isolation [42, 43]. Chromosome fission requires the formation of a new centromere on the newly formed chromosome arm. Generation of synthetic chromosomes by fission has been reported in literature [44, 45]. These observations in the context of muntjac species led us to frame the following question: what is the effect of the condensation of chromosomal content from multiple chromosomes into fewer chromosomes or vice versa on spindle assembly? In other words, what happens to the spindle assembly process when the total chromosomal substance is conserved but re-distributed/partitioned among a varying number of chromosomes? In what follows, we term this conservation of total chromosomal substance as chromosomal volume conservation. Herein, we utilize a general framework of in silico search-and-capture, to determine the effect of chromosomal volume conservation at the initial stages of the spindle assembly process.

Our in silico study indicated that in the absence of chromosomal volume conservation, the total capture time changes monotonically with chromosome number and cell size. However, introducing the total chromosomal volume conservation significantly alters the characteristics of chromosome capture. Our simulations with chromosomal volume conservation further delineated that larger cell size and larger chromosomes promote higher chromosome bi-orientation in the absence of any error correction mechanism. Utilizing this agent-based in silico framework in 3D, we also tested the effect of cell shape on the spindle assembly time and statistics. Changes in cell shape keeping cellular volume conserved were also found to lower the total capture time, when they led to a reduction in the inter-centrosomal distance (pole-to-pole distance noted as centrosome-centrosome distance/CS-CS distance in the rest of the paper), as long as the mobility of the chromosomes was not hindered by the deformations. Slightly oblate cells achieved complete capture sooner than spherical cells whereas prolate cells took longer. Overall, our model alludes that the number of chromosomes prevalent in a species may not be random or accidental. Complex evolutionary mechanisms are likely to be at play to set the number of chromosomes in a species by distributing the total chromosomal content among the chromosomes in a way that favors faithful mitosis.

## II. MODEL SIMULATION

We formulated and employed an agent-based microtubule search-and-capture model based on the literature [4, 10], where microtubules emanating from centrosomes search the cellular volume and capture the chromosomes over the course of time. We simulated our model in 3D geometry by choosing two different shapes for the confinement: (a) spherical and (b) ellipsoidal, depending on the context. In the present model (Fig. 1), chromosomes and kinetochores are modeled as solid 3D cylinders. The sister kinetochores are fixed to the chromosome with their cylindrical axis perpendicular to the axis of the chromosome arm. The size of the kinetochores is kept constant for all simulations (unless specified otherwise), while the length of the chromosome arm is varied. When the total chromosomal volume is mentioned as conserved, we have kept the sum of the arm lengths of individual chromosomes fixed at 20 *μ*m (unless specified otherwise). In one set of simulations, we considered static chromosomes (the chromosomes are fixed in space). In all other cases, we considered diffusive chromosomes where the chromosome movement in space is governed by translational and rotational diffusion.

**FIG. 1.**
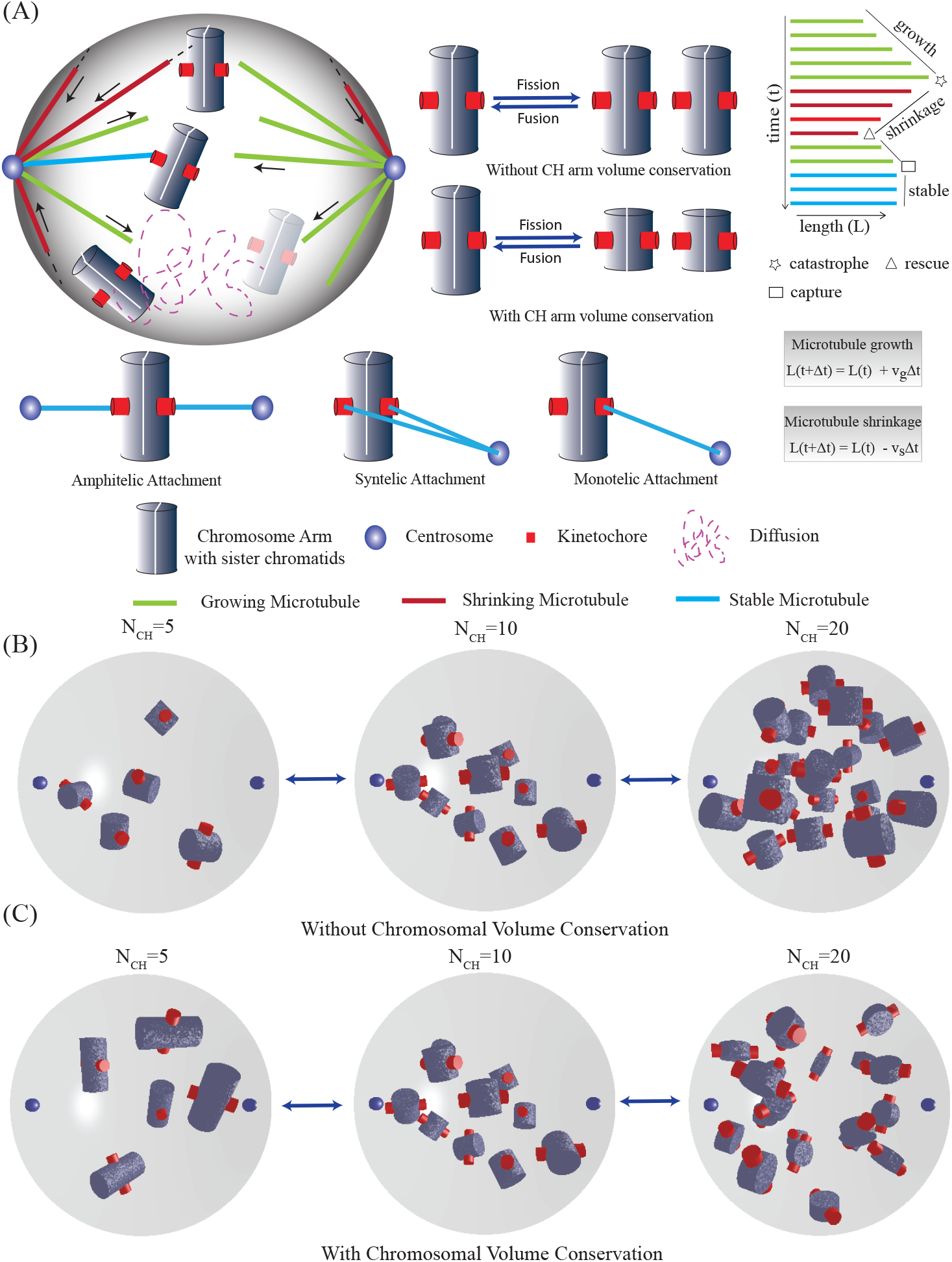
Model schematic and chromosomal distribution. **(A)** Representative illustration depicts various components of the agent-based model. Microtubules (elongating in green, shrinking in red, attached to kinetochore in cyan)) nucleating from centrosomes explore the space to capture sister kinetochores embedded on the chromosome arm. Chromosomal fission and fusion and the effect of chromosomal volume conservation are shown. Microtubules undergo dynamical changes in length while searching for the kinetochores within the cellular volume leading to various types of kinetochore-microtubule attachments. Simulation snapshots for different numbers of chromosomes are shown in **(B)** without chromosome arm volume conservation and in **(C)** with total chromosome arm volume conservation. In the presence of chromosome arm volume conservation, the individual arm lengths of the chromosomes decrease as the number of chromosomes increases. We have chosen the chromosome arm lengths in such a manner that for *N*_*CH*_ = 10 the chromosomes have the same arm length (2 *μ*m) in both (B) and (C).

The centrosomes are placed at the two opposite ends of the diameter (for spherical cells) or a principal axis (for ellipsoidal cells). We modeled microtubules as straight rods of zero thickness emanating from the centrosomes into the 3D cellular volume. The length of the micro-tubule at a particular instant is determined by four dynamic instability parameters (Table II) [4, 6]: growth rate (*v*_*g*_), shrinkage rate (*v*_*s*_), catastrophe frequency (*f*_*c*_) and rescue frequency (*f*_*r*_). Microtubules grow from the centrosomes uniformly in all directions. In the present simulation, no microtubule is rescued after a catastrophe, which means a depolymerizing microtubule shrinks all the way back to the centrosome [10]. A growing microtubule undergoes catastrophe upon hitting (a) the cell periphery or (b) a chromosome arm. A fixed number of microtubules grow from each centrosome. When a growing microtubule runs into a kinetochore, the interaction is stabilized and the event is registered as a capture of the kinetochore. The capture process is completed when all the kinetochores are captured. The biological terms used in this paper and their relevance in the model are summarized in Table I. Model parameters are listed in Table II.

**TABLE I.**
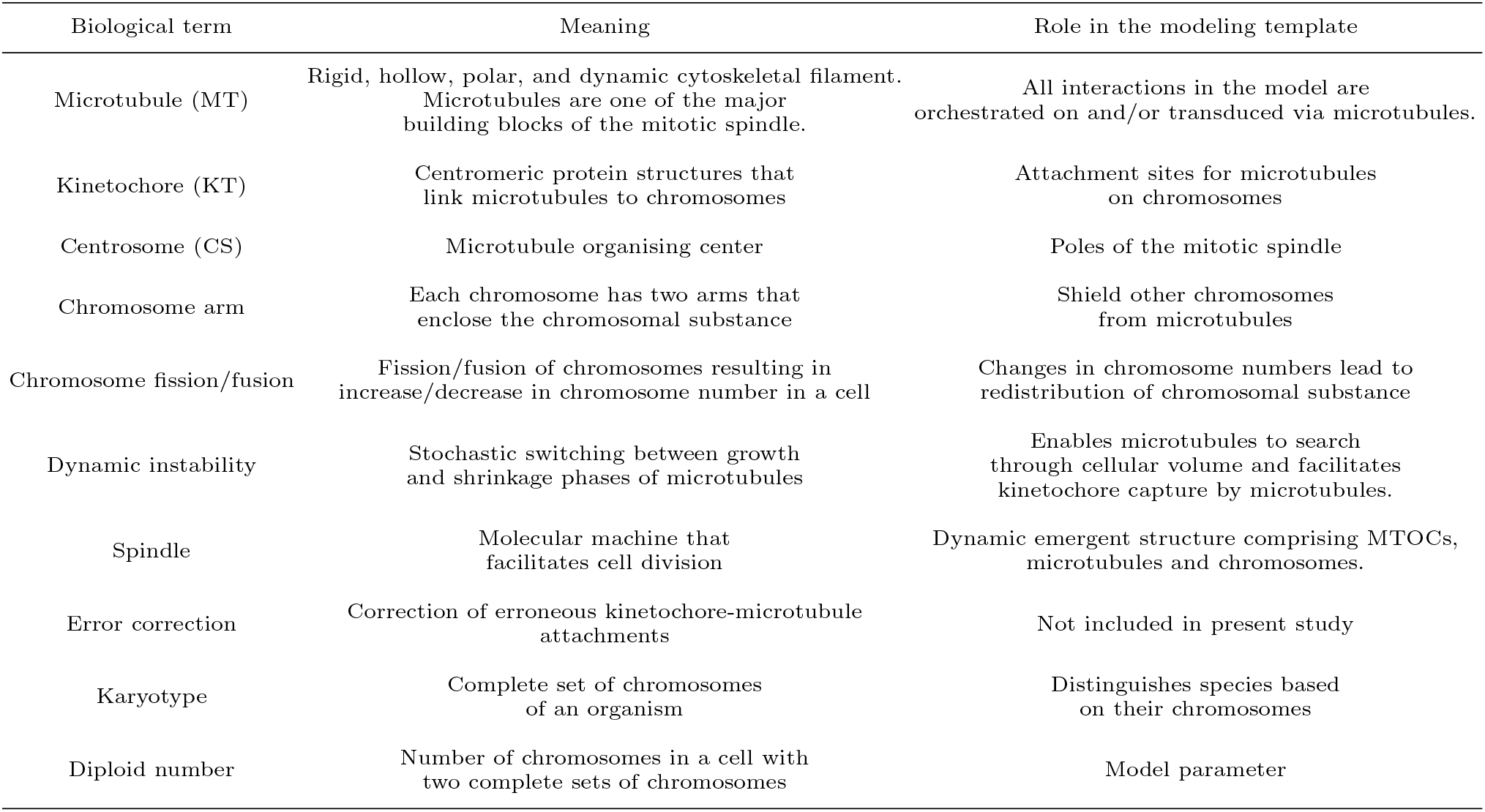
List of biological terms used in this paper and their relevance in the modeling

| Biological term | Meaning | Role in the modeling template |
| --- | --- | --- |
| Microtubule (MT) | Rigid, hollow, polar, and dynamic cytoskeletal filament.<br>Microtubules are one of the major building blocks of the mitotic spindle. | All interactions in the model are orchestrated on and/or transduced via microtubules. |
| Kinetochores (KT) | Centromeric protein structures that link microtubules to chromosomes | Attachment sites for microtubules on chromosomes |
| Centrosome (CS) | Microtubule organising center | Poles of the mitotic spindle |
| Chromosome arm | Each chromosome has two arms that enclose the chromosomal substance | Shield other chromosomes from microtubules |
| Chromosome fission/fusion | Fission/fusion of chromosomes resulting in increase/decrease in chromosome number in a cell | Changes in chromosome numbers lead to redistribution of chromosomal substance |
| Dynamic instability | Stochastic switching between growth and shrinkage phases of microtubules | Enables microtubules to search through cellular volume and facilitates kinetochore capture by microtubules. |
| Spindle | Molecular machine that facilitates cell division | Dynamic emergent structure comprising MTOCs, microtubules and chromosomes. |
| Error correction | Correction of erroneous kinetochore-microtubule attachments | Not included in present study |
| Karyotype | Complete set of chromosomes of an organism | Distinguishes species based on their chromosomes |
| Diploid number | Number of chromosomes in a cell with two complete sets of chromosomes | Model parameter |

**TABLE II.** Parameter values used in the simulations

| Symbol | Description | Value | Reference |
| --- | --- | --- | --- |
| $N_{CH}$ | Number of Chromosomes | 5-40 | This study |
| $R_{cell}$ | Radius of Cell | 8-30 $\mu\text{m}$ | This study |
| $N_{MT}$ | Number of microtubules at each pole | 1000 | [9] |
| $R_{KT}$ | Radius of Kinetochore | 0.44 $\mu\text{m}$ | [10], |
| $R_{CH}$ | Radius of Chromosome | 1.0 $\mu\text{m}$ | [46, 47] |
| $l_{KT}$ | Length of Kinetochore | 0.5 $\mu\text{m}$ | [10] |
| $l_{CH}$ | Length of Chromosome arm | 0.5-4 $\mu\text{m}$ | This study |
| $v_g$ | Microtubule growth rate | 0.35 $\mu\text{m}/\text{sec}$ | [9, 10, 48, 49] |
| $v_s$ | Microtubule shrinkage rate | 1.0 $\mu\text{m}/\text{sec}$ | [9, 10, 48, 49] |
| $f_c$ | Microtubule catastrophe frequency | 0 | This study |
| $f_r$ | Microtubule rescue frequency | 0 | [9, 10, 48, 49] |
| $D$ | Diffusion constant of chromosome | 0.01 $\mu\text{m}^2/\text{sec}$ | [10] |

The code for the agent-based simulation of microtubule-chromosome search-and-capture is written in C. The data analysis and plotting were done in MATLAB (The MathWorks, Natick, MA) and Gnuplot. The computational time for a single simulation run (starting from the onset of microtubule search until all the kinetochores are captured) was in the range of a few minutes (in Intel Xeon CPU having clock speed 2 GHz, RAM 32 GB).

## III. RESULTS

### A. Without chromosomal volume conservation, average capture time changes monotonically with chromosome number

To investigate the dependence of the chromosome search-and-capture process on the chromosomal volume conservation, we first explored how long it takes for the searcher microtubules to capture all the chromosomes without chromosomal volume conservation (Fig. 2A-2B). In this case, the length of each chromosome arm (2 *μ*m) and its volume are fixed. The capture time reported in the following was measured from the time point when the microtubules began to grow from the centrosomes until all the chromosomes were captured. Our results demonstrated that for a fixed radius of the spherical cell, the average capture time increases with increasing chromosome number (Fig. 2A-2B). Clearly, capturing an increased number of targets (chromosomes) by a fixed number of searchers (microtubules) takes longer. Furthermore, we observed that for a fixed number of chromosomes, as the radius of the spherical cell is increased, the average capture time also increases monotonically over a range of cell radii (12 *μ*m -30 *μ*m in the presently explored parameter regime; Fig. 2A-2B). Notice that the chromosomes are distributed randomly within the spherical volume. Therefore, as the cell size increases, the searcher microtubules have to explore a larger volume to find the chromosomes. This contributes to the surge in the average capture time. However, in a small cell (radius ≤10 *μ*m) with a large number of chromosomes (30 ≤*N*_*CH*_ *≤* 40), we observed a noticeable increase in average capture time (Fig. 2B). This deviates from the monotonic trend when the cell radius was varied within 12 *μ*m -30 *μ*m (and 5 ≤ *N*_*CH*_ *≤* 40). Examining the results for a relatively large number of chromosomes (30 *N*_*CH*_ 40) that are ‘tightly’ packed within a relatively small volume (radius ≤ 10 *μ*m), we see that the effective ‘free’ space for the microtubules to ‘pervade’ through and explore is drastically reduced. In a tightly-packed configuration, the chromosome arms occlude the microtubules and induce frequent catastrophe. In addition to that, in crowded confinement, the mobility of the diffusing chromosomes also reduces which in turn negatively impacts the visibility of the kinetochores from the opposite centrosomes. The signature of this specific scenario is evident from the crossover found in Fig. 2B with 30 chromosomes, where the capture time for cell radius 8 *μ*m (red line) increased beyond the capture time for cell radius 12 *μ*m. With 40 chromosomes, the minimum capture time was obtained at a cell radius of 12 *μ*m (Fig. 2B), which is greater than the minimum cell radius explored herein (8 *μ*m). This particular observation alludes that, perhaps for a fixed number of chromosomes ‘tightly’ packed within a small cellular volume, there exists an optimal volume where the average capture time is minimum. Of note, the results presented in Fig. 2A-2B benchmark our computational model setup and are consistent with the previous in silico investigations [10]. We further wanted to check how the capture time varies with chromosome number for static (immobile) chromosomes as a benchmark for our model. However, generating visible configurations (both kinetochores visible to either or both the centrosomes) took enormous computational time as the chromosome numbers (and total volume of chromosomes) were increased. The average capture time of these configurations was also larger than biologically relevant timescales corresponding to mitosis. We refer the interested reader to earlier in-silico works [10] which report that static chromosomes have much larger capture times as compared to mobile chromosomes and the capture time increases exponentially with the number of chromosomes.

**FIG. 2.**
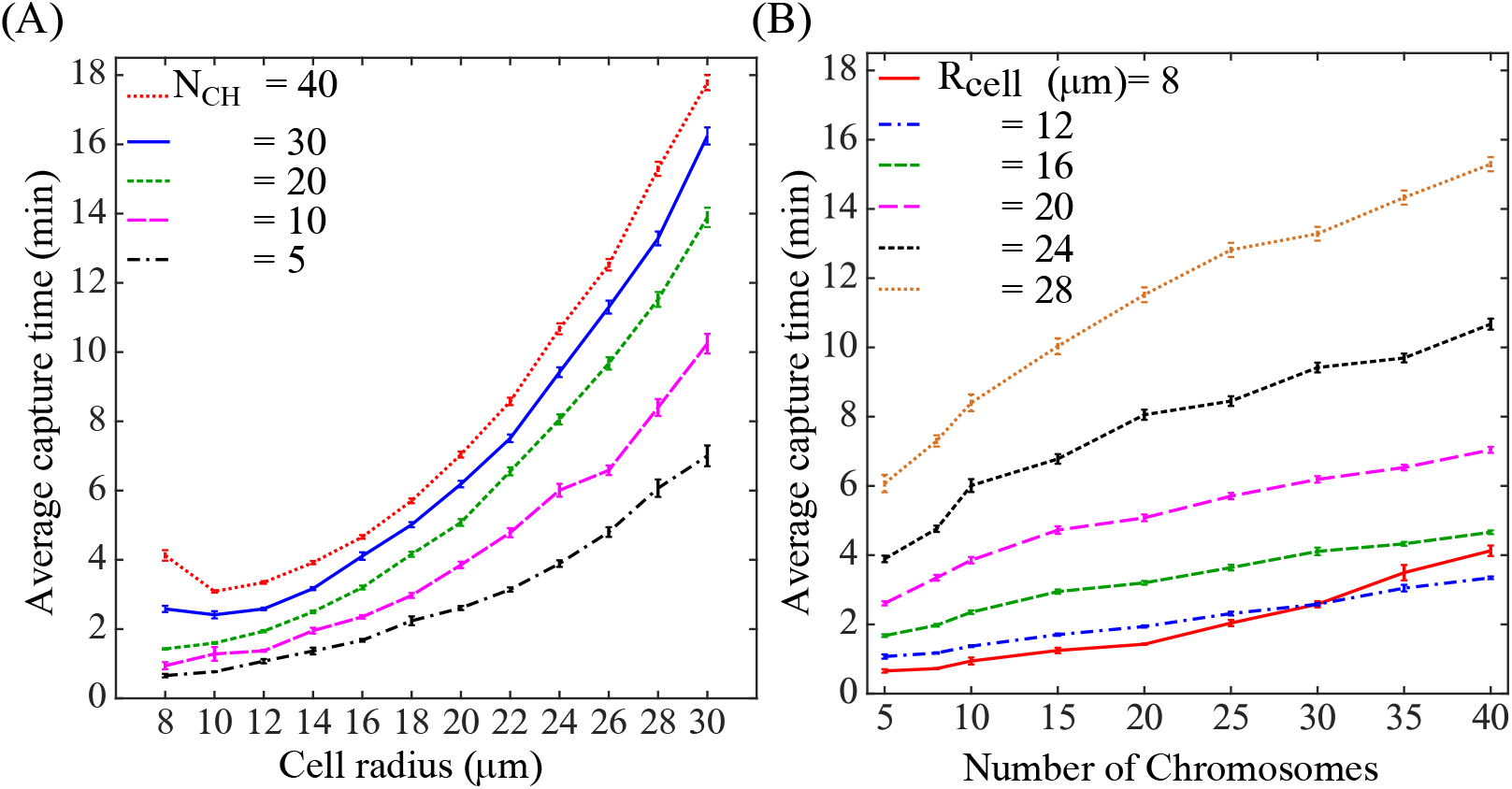
Variation of average capture time with fixed chromosome arm length (2 *μ*m) without chromosomal volume conservation and with mobile chromosomes. **(A)** Capture time as a function of cell radius. Increasing the number of chromosomes hinders chromosomal mobility due to crowding and leads to a larger capture time. For a larger number of chromosomes, an optimal cell radius (e.g. 10 *μ*m for *N*_*CH*_ = 40) is found where the average capture time is the least. **(B)** Capture time increases steadily as a function of the number of chromosomes. The crossover in the capture time for *R*_*cell*_ = 8 *μ*m and 12 *μ*m at chromosome number ∼ 30 indicates the crowding effect in smaller cells. The error bars represent the standard error of the mean.

### B. Conservation of total chromosomal volume significantly alters the characteristics, reduces capture time

Next, we introduced the constraint of total chromosomal volume conservation into our search-and-capture model and investigated the conditions to efficiently capture all the chromosomes. We fix the number of chromosomes to 10 for which the non-conserved and conserved scenarios are identical, i.e. with chromosomal volume conservation, the net volume of 10 chromosomes is equally partitioned among different numbers of chromosomes. We first considered a scenario where the randomly placed chromosomes inside the cell are static. We noticed that the capture time for the chromosomes on increasing their numbers does not increase monotonically to timescales that may be irrelevant to mitosis. Our simulation results showcased an interesting behavior for a fixed cell radius. The average capture time first increased with increasing chromosome number (*N*_*CH*_ *≤* 20) in a fairly monotonic fashion and reached a maximum at *N*_*CH*_ = 20 (Fig. 3A). However, when the number of chromosomes is above 20 (*N*_*CH*_ *≥* 20), with further increase in the number of chromosomes, the average capture time started decreasing and reached a plateau (Fig. 3A). Static Chromosomes showed a maximum capture time when the chromosome arm length equals 1.0 *μ*m (Fig. 3A). According to the model, with the increase in chromosome number, the individual chromosome arm length was decreased to conserve the total chromosomal volume (Fig. 1). The initial increase in the average capture time with increasing chromosome number (up to *N*_*CH*_ = 20) is due to the increase in the number of targets (chromosomes), while the number of searchers (microtubules) remained fixed. This is further aided by the fact that, for static chromosomes, the number of chromosome arm-induced microtubule catastrophes decreases with decreasing individual chromosome length. In this regard, note that a premature microtubule catastrophe during an unsuccessful search (e.g., hitting a large chromosome arm) enhances the chance of a new microtubule cycle which in turn may increase the efficiency of the search and capture. Now, when the chromosome number *N*_*CH*_ is above 20, the chromosome arm length falls below 1 *μ*m due to volume constraint (Fig. 3A). In this scenario, the size of the kinetochores (∼ 0.44 *μ*m) becomes comparable to the chromosome arm length. According to the model construction, when the chromosome arm length becomes shorter than the kinetochore, the chromosome arm is only partially ‘exposed’ and most of it is ‘buried’ within the kinetochore. This results in a drastic reduction in the number of chromosome-arm-induced microtubule catastrophes, which, however, enhances the visibility of sister kinetochores from the opposite centrosomes. The enhanced visibility reduces the average capture time significantly with increasing chromosome numbers even though the number of microtubules remained fixed. The average capture time reaches a plateau at *N*_*CH*_ *≥* 35 in the currently explored parameter regime (Fig. 3A) when there is no effect of the chromosome arm. The scenario is comparable to chromosomes that are transparent to the microtubules [9].

**FIG. 3.**
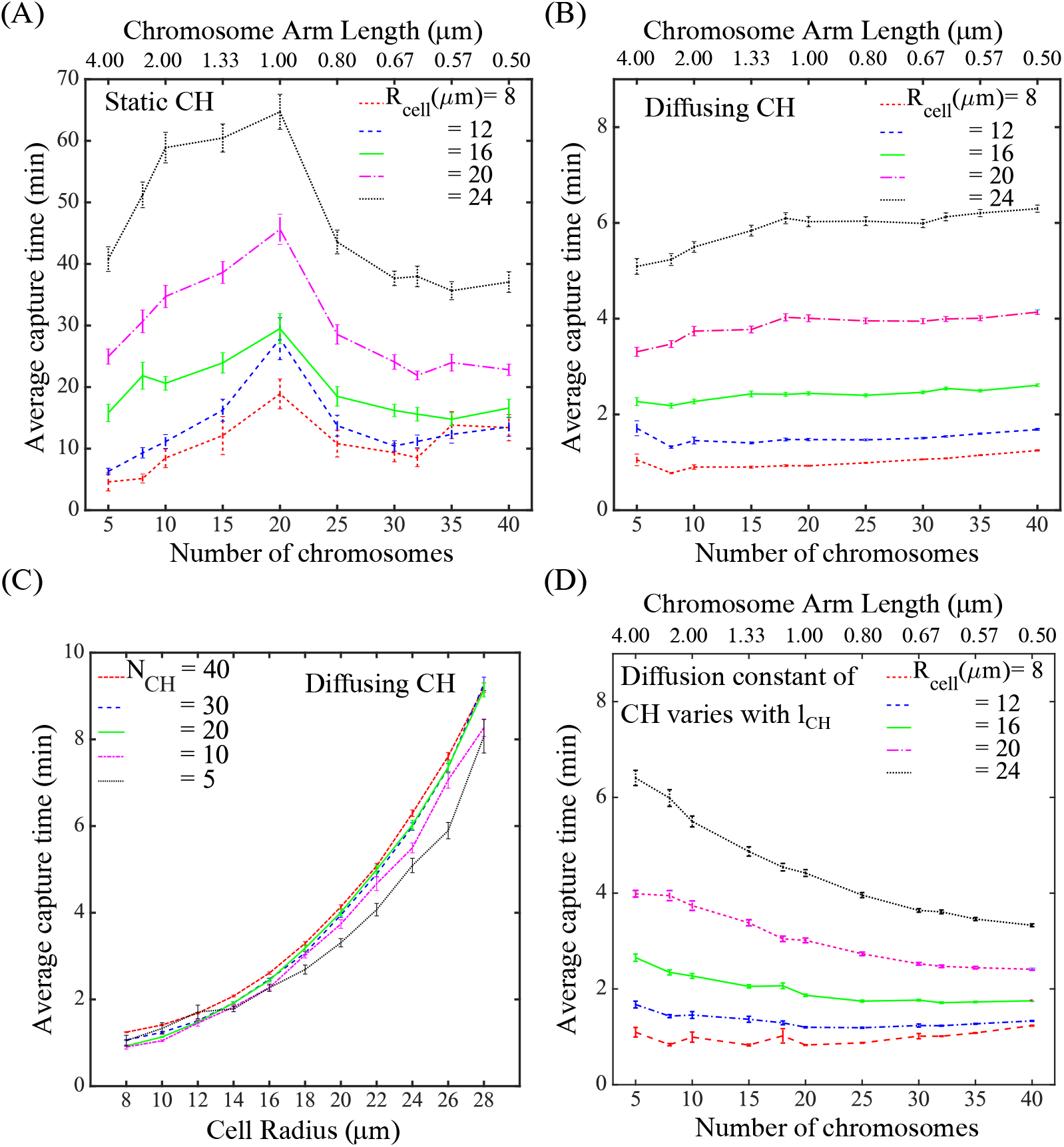
Variation of average capture time with chromosomal volume conservation. **(A)** For static chromosomes, *Capture time* increases steadily with *chromosome number* up to *N*_*CH*_ = 20 and then decreases sharply before saturation. Total chromosome arm length is conserved at 20 *μ*m. Configurations with all kinetochores visible to microtubules from either or both the centrosomes were chosen for the simulations. **(B)** *Capture time* is substantially reduced and varies marginally with *chromosome number* for *diffusing* chromosomes, compared to the static chromosomes in (A). Diffusing chromosomes have much smaller average capture times compared to static chromosomes. **(C)** *Capture time* increases monotonically with *cell radius* without much variation for different *N*_*CH*_. No increase in capture time is observed due to the crowding of chromosomes even at small cell sizes (compare Fig. 2A). **(D)**Variation of capture time with chromosome number keeping diffusion constant 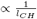 such that the diffusion constant for chromosome arm length 2.0 *μ*m is 0.01 *μ*m^2^/sec. A larger chromosome arm refers to higher viscous drag and less mobility of chromosomes thereby increasing the capture time.

Next, we explored the effect of the chromosomal volume conservation on the search-and-capture process when the chromosomes, instead of being static, diffuse inside the cell. We found that the diffusing chromosomes are captured much faster than static chromosomes (Fig. 3B). The average capture time, for diffusing chromosomes, increased only slightly with chromosome numbers for cell radius exceeding 20 *μ*m. For cell radius less than 20 *μ*m, capture time was almost constant for all chromosome numbers (Fig. 3B). However, a slight increase in the capture time was observed in small cells with radius 8 *μ*m and 12 *μ*m for fewer chromosomes (i.e., when the chromosome arm lengths are large ∼.0*μ*m). This suggests large chromosome arms may reduce the mobility of chromosomes in constricted cells as well as reduce the visibility of the sister kinetochores leading to delayed capture. Additionally, we observed that the average capture time is less sensitive to chromosome number variation in the presence of chromosome arm volume conservation (compare Fig.2A and Fig. 3C). Exploring the chromosomal configurations, we see that as the chromosome number is increased (in Fig. 3C), chromosome arms become shorter to maintain the volume conservation. As a result, the effect of steric hindrance on the chromosomes is significantly reduced. Shorter chromosome arms increase the overall visibility of the chromosomes across a range of chromosome numbers in the explored parameter regime. We also checked how the capture time varies with chromosome numbers when the diffusion constant *D* is scaled as the inverse of chromosome arm length 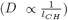. The underlying rationale behind this variation is that the larger chromosomes are expected to have lesser mobility. The diffusion constant remains unaltered when the number of chromosomes *N*_*CH*_ = 10. Our results show that in large cells having fewer chromosomes with long chromosome arms, the capture time increases significantly (see Fig. 3D). In smaller cells, the increase in the capture time for fewer chromosomes is only marginal. Reduction of mobility due to higher viscous drag applied on the larger chromosomes plays a key role in increasing the capture time in larger cells. However, this effect does not contribute much in smaller cells where the crowding of the chromosomes and restricted orientations are major reasons for the reduced mobility [50].

### C. Larger cells and larger chromosomes lead to higher chromosome bi-orientation

Next, we ask about the fidelity of the chromosomal segregation in the context of chromosomal volume conservation. We acknowledge that for proper spindle assembly, search-and-capture is a necessary but not a sufficient condition that ensures chromosomal biorientation. In addition, it is crucial to have the correct type of attachment between the centrosomes and chromosomes for faithful cell division. The microtubule-chromosome attachments can be classified into the following categories depending on their connection with the centrosomes: (a) amphitelic, (b) monotelic, (c) syntelic, and (d) merotelic (Fig. 1). The stability of these attachments differs from one another- the amphitelic attachments are the most stable, ‘correct’ type of attachment that facilitates proper chromosomal biorientation and faithful segregation. On the other hand, the non-amphitelic attachments are erroneous and promote aneuploidy. During the spindle assembly process, several ‘error correction’ mechanisms specifically destabilize the erroneous attachments so that only the correct amphitelic attachments prevail (recently reviewed in [51]). Although our present model does not include the correction of erroneous attachments, we asked the following question: without the intervention of any error correction mechanism, what is the relative fraction of amphitelic attachments, when all the chromosomes are captured? We further wondered-how important the positioning of the chromosomes are when the amphitelic attachment is formed. Is there a preferential location inside the cell where one type of attachment is formed more frequently compared to another? To address this, we tracked individual chromosomes in the simulation and measured how far the chromosomes were from the *equatorial* plane when both the sister kinetochores were captured by microtubules from *left* and *right* centrosomes (Fig. 4A). We found that the chromosomes that got captured near the centrosomes were mostly syntelic attachments (sister kinetochores captured from the same pole) (refer to Fig. 8A). Amphitelic attachments, espe-cially the capture of the uncaught kinetochore (second attachment in Fig. 4A), mostly occurred when the chromosomes were in the vicinity of the equator (Fig. 4B, Fig. 8E-8F). This suggests that the chromosomes near the metaphase plate have higher chances of establishing amphitelic attachments (in the absence of any error correction) compared to the chromosomes that are close to the poles. For smaller cells (e.g. *R*_*cell*_ = 8 *μ*m), amphitelic capture distribution had a sharp peak near the central region whereas the distribution peak decreases for larger cell radii (Fig. 4B). There exists a slice of volume within the cell around the metaphase plate where the chance of a chromosome capture being amphitelic is the highest. This can also be justified by the fact that amhitelic attachments start forming after an initial delay from the start of the simulations which can be attributed to the time taken by the searcher microtubules from both centrosomes to reach the centrally situated chromosomes (Fig. 8D). The results qualitatively remained the same when the constraint of chromosomal volume conservation was lifted (Fig. 4C). We next explored how the probability of amphitelic attachments depends on the size of the spherical cell and the number of chromosomes both in the presence and absence of chromosomal volume conservation. We found that in the presence of chromosomal volume conservation, the population of amphitelic attachments grows as the radius of the cell increases (Fig. 4D). This behavior fairly holds true for almost the entire range of chromosome numbers scanned herein (*N*_*CH*_ *∼* 5 -40; Fig. 4D). We further observed that for a fixed cell radius, the probability of having amphitelic attachments is slightly higher for a smaller number of chromosomes (e.g., in Fig. 4D, for cell radius 20 *μ*m, compare *N*_*CH*_ = 10 and 40). This alludes that chromosomes with larger arms (i.e., for a small number of chromosomes, in the presence of chromosomal volume conservation) are likely to form more amphitelic attachments with microtubules from opposite centrosomes. Larger chromosome arms reduce the visibility of the distal sister kinetochore from the capturing centrosome thereby reducing the probability of a syntelic attachment. Furthermore, if the entire chromosomal substance is distributed among fewer chromosomes, the number of attempts needed for capturing all the chromosomes is also less. This reduces the possibility of wrong attachments and increases the relative fraction of amphitelic attachments. However, in cells with smaller radii, the chromosomes are congested and the effect of steric hindrance is considerable. In that scenario, chromosomes that have already established a monotelic attachment find it difficult to diffuse toward the opposite pole. Being *entrapped* near the capturing pole increases the chance of the sister kinetochore being captured by the same pole leading to the syntelic attachment. Amphitelic attachments in general increase with cell size (Fig. 4D-4E). As the cell size increases, the chromosomes find more space to diffuse and can move toward the opposite pole to be captured and establish amphitelic attachments.

**FIG. 4.**
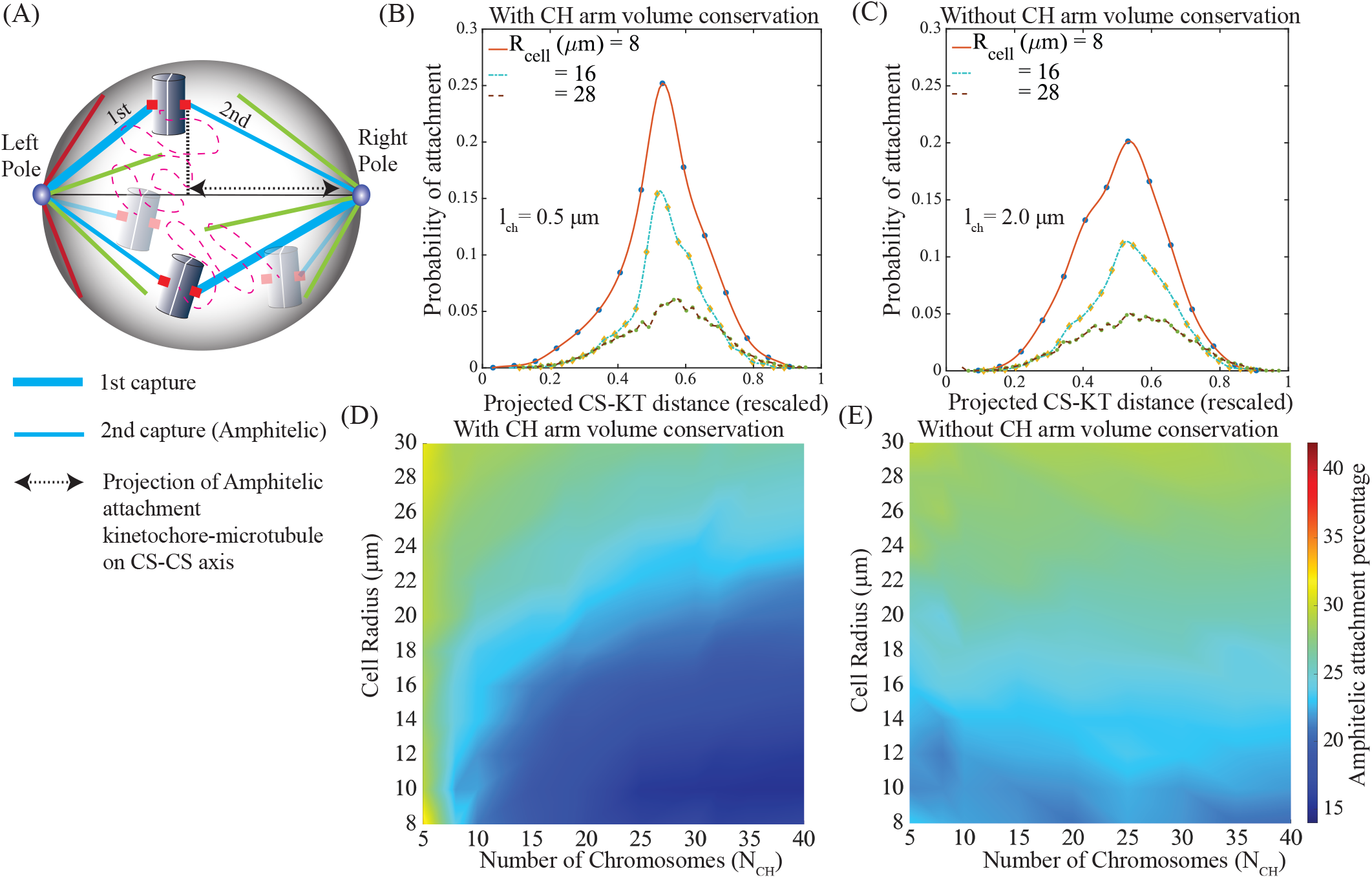
Statistics of correct spindle for different cell sizes. **(A)** Schemes for sequential capture of sister kinetochores leading to amphitelic attachment. Centrosome (CS)-kinetochore (KT) capture distance is measured by taking projections of the microtubule length on the centrosome-centrosome axis. **(B)** Distribution of amphitelic capture distance (2*nd* capture) from capturing centrosome for 40 chromosomes (distances were rescaled by cell diameter). Total chromosome arm length was conserved at 20 *μ*m (individual arm length was 0.5 *μ*m). **(C)** Distribution of amphitelic capture distance (distances were rescaled by cell diameter) from capturing centrosome without chromosomal arm volume conservation for 40 chromosomes (individual arm length was 2.0 *μ*m). In both (B) and (C) the distributions have sharp peaks for small cell radii which are absent in large cells. The location of the peaks suggests that the amphitelic capture mostly happens in the central region. **(D)** Percentage of amphitelic attachments among chromosomes (CH) with conserved chromosome arm volume. A higher percentage of amphitelic attachment was observed for small *N*_*CH*_ with chromosomal volume conservation. **(E)** Percentage of amphitelic attachments among chromosomes without chromosome arm volume conservation. In larger cells, amphitelic percentage remains high across the range of *N*_*CH*_. Color bar represents percentage of amphitelic attachments among chromosomes.

### D. Effect of cell shape on the spindle assembly time and statistics

Next, we tested how the chromosomal capture depends on the shape of the cellular confinement (Fig. 5) in the presence of total chromosomal volume conservation. We started with a spherical volume and systematically changed the length of one of the axes (the axis which joins the opposite centrosomes), while the other two axes were kept equal in a manner such that the total volume was constant. We found that the average capture time is higher in prolate-shaped cells (Fig. 5A).

**FIG. 5.**
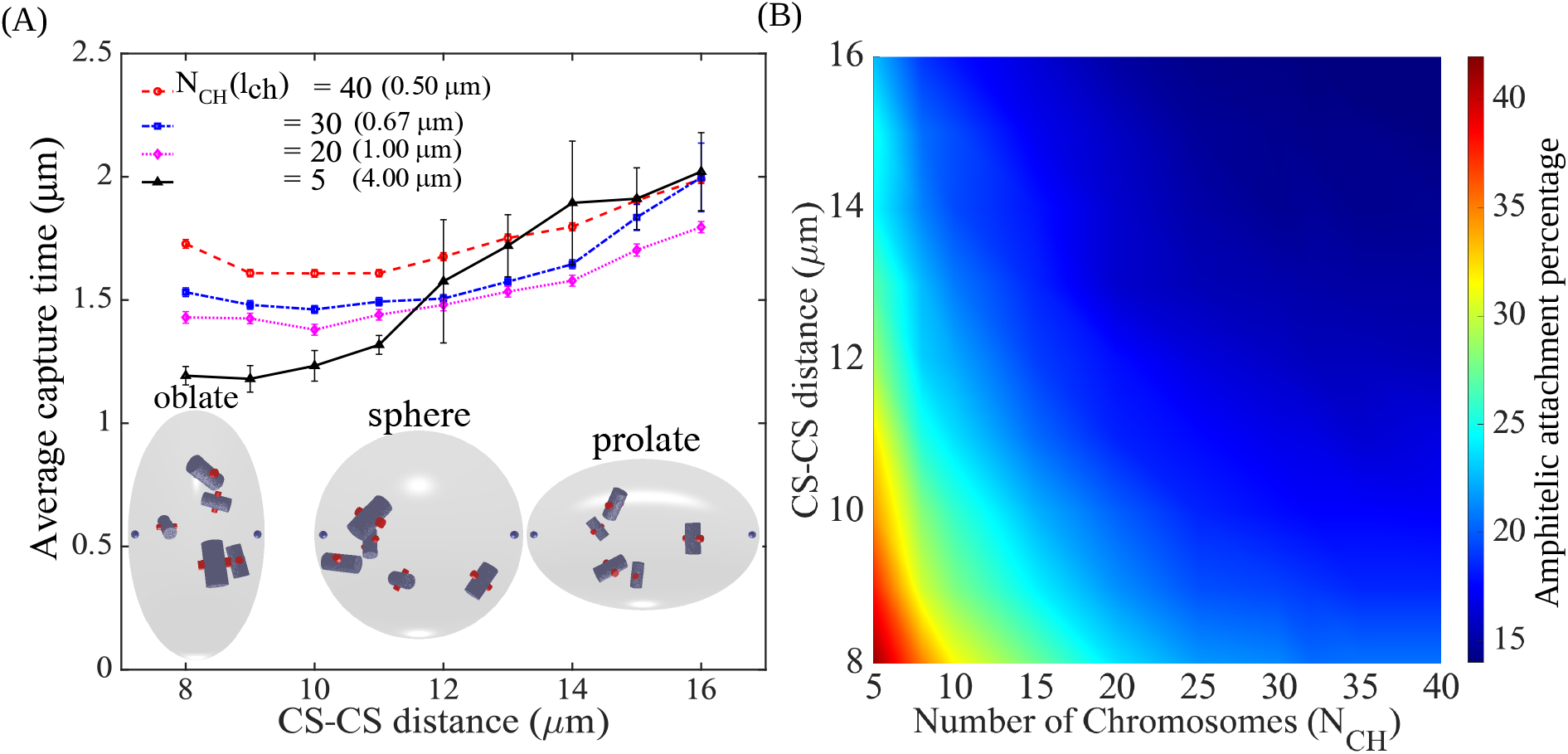
Statistics of correct spindle for different cell shapes. (A) Variation of *capture time* with *centrosome-centrosome distance* in presence of total chromosome arm length (and volume) conservation. The chromosome arm lengths are indicated next to the chromosome number. Rapid increase in the average capture time for *N*_*CH*_ = 5 for larger centrosome-centrosome distance is due to long chromosome arm (4 *μ*m) that occludes microtubules. (B) Amphitelic attachment percentage as a function of chromosome number and centrosome-centrosome distance. The total chromosome arm length was conserved at 20 *μ*m. The cell is spherical for centrosome-centrosome axis length 12 *μ*m, oblate with the centrosomes placed at the shorter axis for centrosome-centrosome axis length *<* 12 *μ*m, prolate with the centrosomes placed at the larger axis for centrosome-centrosome axis length *>* 12 *μ*m. The color bar indicates the percentage of amphitelic capture. Searcher microtubules in prolate cells have to travel longer to reach near the metaphase plate where most of the amphitelic attachments happen. This reduces the probability of amphitelic attachment.

Within a prolate volume (intercentrosomal distance *>* 12 *μ*m in Fig. 5A), the searcher microtubules have to travel a longer distance to capture chromosomes situated closer to the metaphase plate compared to spherical volume. This slightly increases the capture time. We also found that in oblate volume (intercentrosomal distance *<* 12 *μ*m in Fig. 5A), if the eccentricity is too high the capture time increases slightly compared to the spherical volume. This marginal increase (*N*_*ch*_ = 40, dashed red curve in Fig. 5A) may be attributed to the fact that chromosomes situated near the rim of the metaphase plate in an oblate volume are farther away from the centrosomes than in spherical volumes. Thus microtubules have to travel longer distances to capture them. However, for *N*_*ch*_ = 5, shorter intercentrosomal distances in oblate volumes aid toward reduced capture time compared to spherical volumes (Fig. 5A), solid black curve). We also found that chromosomes experience more boundary collisions in prolate volume as compared to spherical and oblate volumes (refer to inset in Fig. 8C). This may lead to chromosomes with larger arms experiencing reduced mobility in a prolate volume and as a result, the average capture time is higher (Fig. 5A, solid black curve).

We also checked the relative fraction of amphitelic attachment as a function of the cell shape (Fig. 5B). Our simulations demonstrated that the relative fraction of amphitelic attachments is lower in prolate volumes (intercentrosomal distance *>* 12 *μ*m in Fig. 5) compared to spherical and oblate volumes (intercentrosomal distance ≤12 *μ*m in Fig. 5) when the number of chromosomes *N*_*CH*_ is below 20 (Fig. 5B). Note that, in prolate cells, most of the chromosomes need to travel a larger distance to become accessible to microtubules from opposite centrosomes. This condition promotes amphitelic attachment. However, once a chromosome is partially captured near the centrosome, it diffuses around but cannot move swiftly toward the opposite centrosome. Consequently, both the sister kinetochores are captured from the same pole, leading to the syntelic attachment. On the other hand, in oblate cells (shorter CS-CS distance), sister kinetochores encounter microtubules at similar probabilities from the opposite poles. This, in turn, facilitates frequent amphitelic attachments. Consistent with the previous trend as shown in Fig. 4D), in this case too, the formation of amphitelic attachments is more common when the chromosome arms are larger (in other words, the number of chromosomes is small as the constraint of chromosomal volume conservation is in place). Of note, the dependence of spindle assembly speed and accuracy on the structural and mechanical parameters of the chromosomes was recently investigated in [36, 50] utilizing similar 3D agent-based computational models.

## IV. DISCUSSION

The formation of a proper mitotic spindle is essential for faithful mitosis. Karyotype variations can affect the process of mitosis significantly. The survival of an organism with a specific karyotype depends greatly upon its ability to successfully perform mitotic cell division. Changes in chromosome number can happen due to inter-species mating or genomic mutations that generally produce infertile offspring due to a mismatch of cell numbers between the parent cells and the daughter cell [41, 52–54]. It is hypothesized that whole genome duplication events followed by chromosomal rearrangements [55] in such daughter cells is an evolutionary process to restore fertility [52, 56–59]. Whole genome duplication leads to a doubling of the chromosome number in a cell. Further, chromosome fission/fusion in the parent cell can also occur due to mutations as often found in cancer cells [60–62]. Cancer cells are also prone to whole genome duplication events [60–62]. We checked how such changes in the chromosome number affect the mitotic spindle assembly. Our study elucidates that the altered karyotype in the daughter cell has a lower average capture time when the total number of chromosomes decreases and the chromosomal volume is not conserved. An increase in chromosome numbers without chromosome arm volume conservation can lead to larger capture times [60]. A decrease in net cell volume can reduce the capture time significantly. If the total chromosomal volume is not conserved, decreasing cell volume can lead to crowding and higher average capture time in cells with many chromosomes. However, if the total chromosomal volume is conserved, the cap-ture time roughly remains the same for all chromosome numbers. When Cytoplasmic viscous forces are applied proportional to the size of the chromosome arm, the mobility of larger chromosomes reduces. This effectively increases the capture time.

Increasing chromosome numbers with total chromosomal volume conserved reduces the percentage of amphitelic attachments as the size of the chromosome arm becomes smaller. However, amphitelic attachment percentage does not vary much with chromosome numbers when the total chromosomal volume is non-conserved (Fig. 6). We also found that higher cell volume favors amphitelic attachments due to less crowding of the chromosomes [50]. Prolate cell geometry reduced the possibility of amphitelic attachments whereas oblate cells with the same total volume showed a higher probability of amphitelic attachment. This occurs because microtubules have to travel a longer distance to reach the chromosomes situated near the metaphase plane in prolate confinements as compared to oblate confinements (see Fig. 6). Telocentric chromosomes were found to have lower capture times for long chromosomes compared to metacentric chromosomes. Shielded telocentric chromosomes can become visible by rotational diffusion only, however, metacentric chromosomes require translational diffusion to become completely visible (see appendix Fig. 9B). If the total chromosomal volume is randomly distributed among the chromosomes, the presence of some large chromosomes can increase the capture time. However, the capture time statistics remain qualitatively similar to the scenario when the total chromosomal volume is uniformly distributed (see appendix Fig. 9D).

**FIG. 6.**
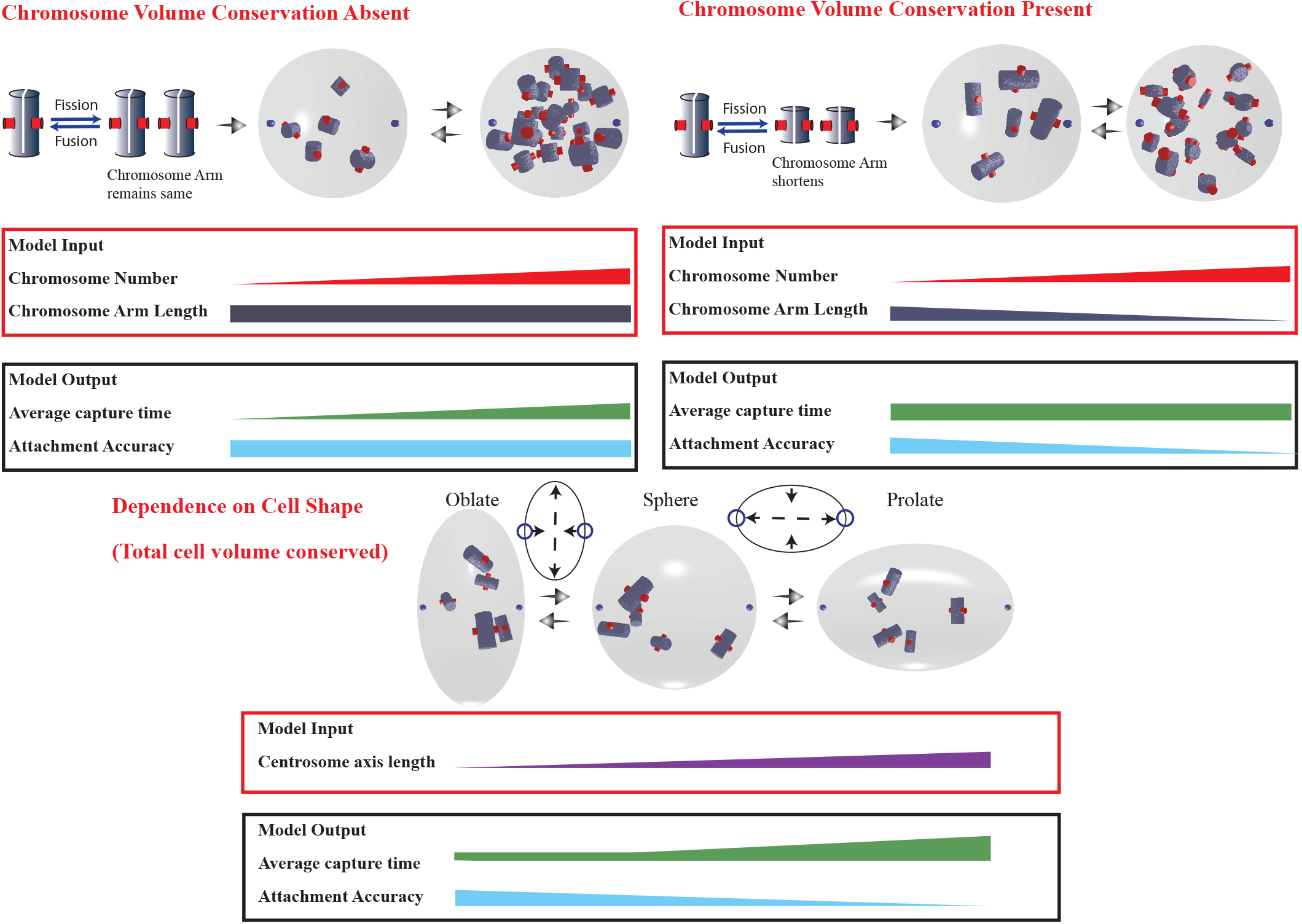
Graphical summary of model outcomes outlining the dependence of microtubule-chromosome search and capture on chromosomal volume conservation. According to our agent-based simulations, in the absence of chromosomal volume conservation, although the average capture time increases with increasing chromosome number, the attachment accuracy (in other words, the probability of having correct amphitelic attachments) remains unchanged. In the presence of chromosomal volume conservation, the average capture time does not change with the increase in chromosome numbers but the accuracy of kinetochore-microtubule attachments decreases. When the cell shape is varied keeping the total cell volume conserved, capture time increases slightly with the increase in intercentrosomal distance and kinetochore-microtubule attachment accuracy decreases.

Needless to say that changes in chromosome numbers in a cell may or may not be accompanied by the conservation of total chromosomal volume. Chromosome number reduction in yeast strains reportedly follows a pathway of a centromere deletion from one chromosome and then fusion of the rest of the chromosome to another that contains a working centromere [63]. Our study elucidates that under chromosomal volume conservation, a fusion of chromosomes accelerates the process of mitotic spindle assembly by favoring correct attachments (see Fig. 6). However, processes such as whole genome duplication are bound to increase the total capture time and delay mitotic spindle assembly as the net chromosomal volume is not known to be conserved in such processes [60].

This study is primarily focused on the effects of total chromosomal volume conservation on the spindle assembly time and the nature of microtubule-kinetochore attachments.Our present simplistic model does not include molecular motors like dynein and kinesin and the impact of the forces stemming from their mechanochemical activity on the dynamics of mitotic spindle [64, 65]. The centrosomes in our model are static and there is no additional mechanism that corrects erroneous attachments [66, 67]. It would be a viable future study to consider the effects of various molecular motor-mediated forces on the mitotic spindle and erroneous attachment correction mechanism in the presence of chromosome arm volume conservation.

## V. ACKNOWLEDGMENTS

R.P. acknowledges the fellowship from SERB (Science and Engineering Research Board), Department of Science and Technology (DST), India (EMR/2017/001346). P.N. was supported by a fellowship from CSIR, India.

R.P. conceived and directed the study. S.C., P.N. and R.P. wrote the manuscript. S.C. and P.N. performed the theoretical modeling and contributed equally to this paper. All coauthors contributed to the data analysis, and editing of the manuscript and approved the content.

## Appendix A: Supporting data

### 1. Chromosome arm induced catastrophe increases with increasing chromosome arm length

To ensure the correctness of our inference that larger chromosome arms lead to amphitelic captures, we checked the number of chromosome arm-induced microtubule catastrophes under various conditions for both static and mobile chromosomes. Our results showed that mobile chromosomes induced much more microtubule catastrophes than static ones as their chances of encountering a microtubule tip are higher (Fig. 7). The chromosomes with larger arms also induced more catastrophes than the ones with smaller arms. For static chromosomes, if the total chromosomal volume is conserved, increasing the number of chromosomes resulted in little increase in the number of arm-induced catastrophes. However, if chromosome numbers are increased without conserving the total chromosomal volume, arm-induced catastrophe increases manifold (Fig. 7). A similar trend is also seen in mobile chromosomes, but even with chromosome arm volume conservation, increasing chromosome numbers greatly increased the number of arm-induced catastrophes (Fig. 7). Overall, a larger chromosome arm leads to more shielding of opposite-facing kinetochore from one pole and thereby amphitelic attachment becomes more probable.

**FIG. 7.**
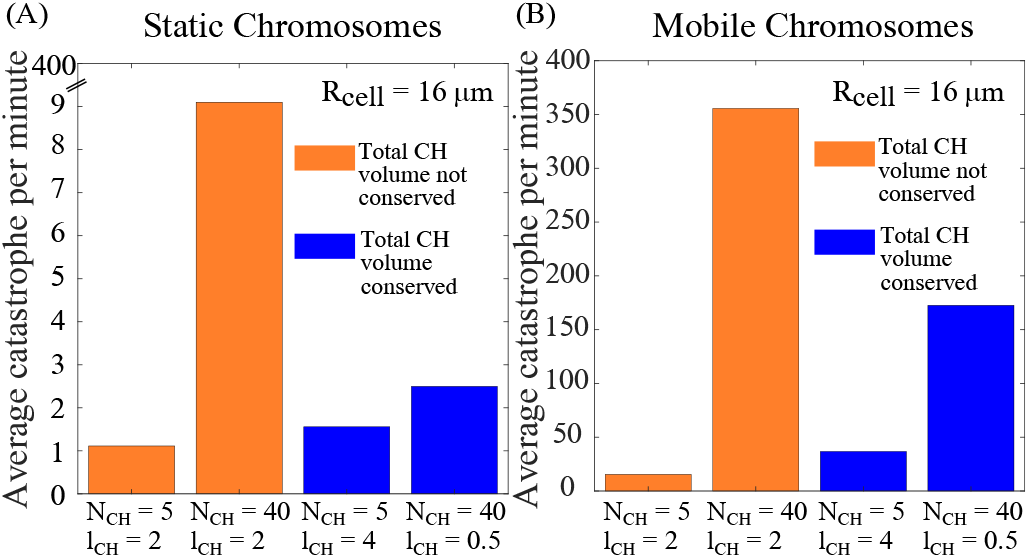
Comparison of chromosome arm induced microtubule catastrophe per minute for static (A) and mobile chromosomes (B). Yellow bars indicate the data without chromosome arm volume conservation and blue bars with chromosome arm volume conservation. Statistical average was taken over a hundred ensembles. All simulations were run for 10 minutes of simulation time.

### 2. Syntelic captures mostly occur near the poles

Chromosomes situated near the pole find more microtubules from the closer pole. Hence, they have a higher probability to form syntelic attachments. Microtubules from the distant pole have to travel a larger distance to capture these chromosomes and mostly they undergo catastrophe before reaching these chromosomes and forming a kinetochore-microtubule attachment. Our data clearly show that the peak of the syntelic distribution is located near the poles (Fig. 8A). The peak of the syntelic distribution is located between the poles and the metaphase plate. This happens because the steric effects of the chromosomes prevent too many chromosomes from crowding near the poles(Fig. 8A (orange line)).

**FIG. 8.**
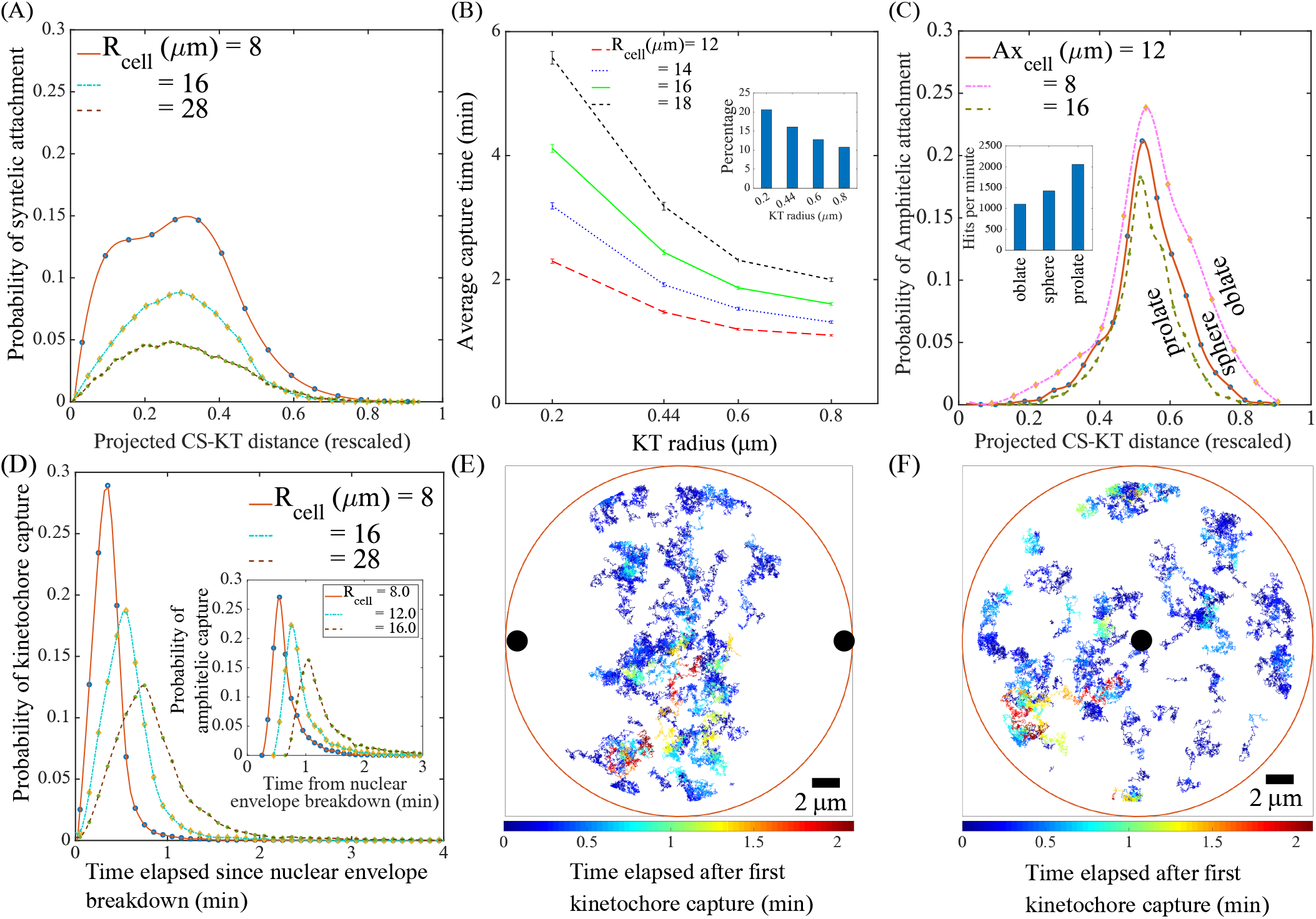
**(A)** Distribution of projected syntelic capture distance from capturing centrosome with distances rescaled by cell diameter. Distribution shown for 40 chromosomes (chromosome arm length 0.5 *μ*m). **(B)** Variation of capture time with kinetochore radius. **Inset** shows percentage of amphitelic attachments for *R*_*cell*_ = 12.0 *μ*m and various kinetochore radii. Each data set contained 20 chromosomes with chromosome arm length 1.0 *μ*m. **(C)** Distribution of projected amphitelic capture distance from capturing centrosome with distances rescaled by centrosome-centrosome axis length. Data set contained 40 chromosomes (chromosome arm length 0.5 *μ*m). **Inset** shows the number of boundary collisions per minute of 5 chromosomes (chromosome arm length 4.0 *μ*m) in various cell shapes. **(D)** Temporal evolution of probability of kinetochore capture. **Inset** shows temporal evolution of amphitelic captures. Data set contained 40 chromosomes (chromosome arm length 0.5 *μ*m) (**E**) Projection of chromosome trajectory till amphitelic capture on plane containing both centrosomes. **(F)** Projection of chromosome trajectory till amphitelic capture on metaphase plate. Temporal evolution of chromosomes in (E),(F) is shown from capture of first sister kinetochore till amphitelic capture; black spheres represent centrosomes.

### 3. Larger kinetochore leads to faster capture but increases erroneous captures

We also explored the effects of a larger kinetochore size on the average capture time. Our results (Fig. 8B) showed that the average capture time decreases with larger kinetochore size but amphitelic attachments decrease. A larger kinetochore, therefore, helps in faster capture of the chromosomes but at the same time, the chances of erroneous attachments increase [50]. Thus, even if there is accelerated capture due to a larger kinetochore size, there would be more erroneous microtubule-kinetochore attachments to correct. Earlier computational work [50] suggests an optimized kinetochore surface area exists that minimizes the capture time as well as reduces the number of erroneous attachments.

### 4. Cell shapes may affect the mobility of chromosomes and reduce bi-orientation

For all variations in cellular confinement studied, the chromosomes that attain amphitelic attachment are mostly situated near the metaphase plate as shown in Fig. 8C. The searcher microtubules have to travel longer to reach the central region in prolate confinement compared to oblate and spherical confinements. Furthermore, chromosomes in oblate cells experience much fewer boundary collisions compared to chromosomes in prolate cells. This contributes to the reduction of their mobility. These factors may play a leading role in decreasing the number of amphitelic attachments in prolate cells.

### 5. Telocentric chromosomes have lower capture time when chromosome numbers are low

Telocentric chromosomes have the kinetochores located at one end of the chromosome arm (Fig. 9A). Due to this configuration, a telocentric chromosome within a chromosomal crowd needs only rotational diffusion to expose its kinetochores to the centrosomal microtubules. Under a similar situation, kinetochores of a metacentric chromosome would not become visible only by such rotations. Apart from rotational diffusion, translational diffusion is needed to make the shielded metacentric chromosomes visible to the microtubules. This effect however is predominant only when the number of chromosomes is small (see Fig. 9B, for 5-10 chromosomes). We ran simulations by setting the translational diffusion coefficient to zero for both telocentric and metacentric chromosomes in the large chromosome arm regime. The results show that with only rotational diffusion present, telocentric chromosomes have a lower capture time than metacentric chromosomes (see Fig. 9B (Inset)). In this regime, with the increasing number of chromosomes, capture time increases and becomes nearly the same as that of metacentric chromosomes. Further increase in the chromosome number severely shortens the chromosome arms and the kinetochores become completely visible irrespective of telocentric and metacentric chromosomes (compare Fig. 9B with Fig.3D). For short chromosomes the capture time varies similarly to Fig. 3D.

**FIG. 9.**
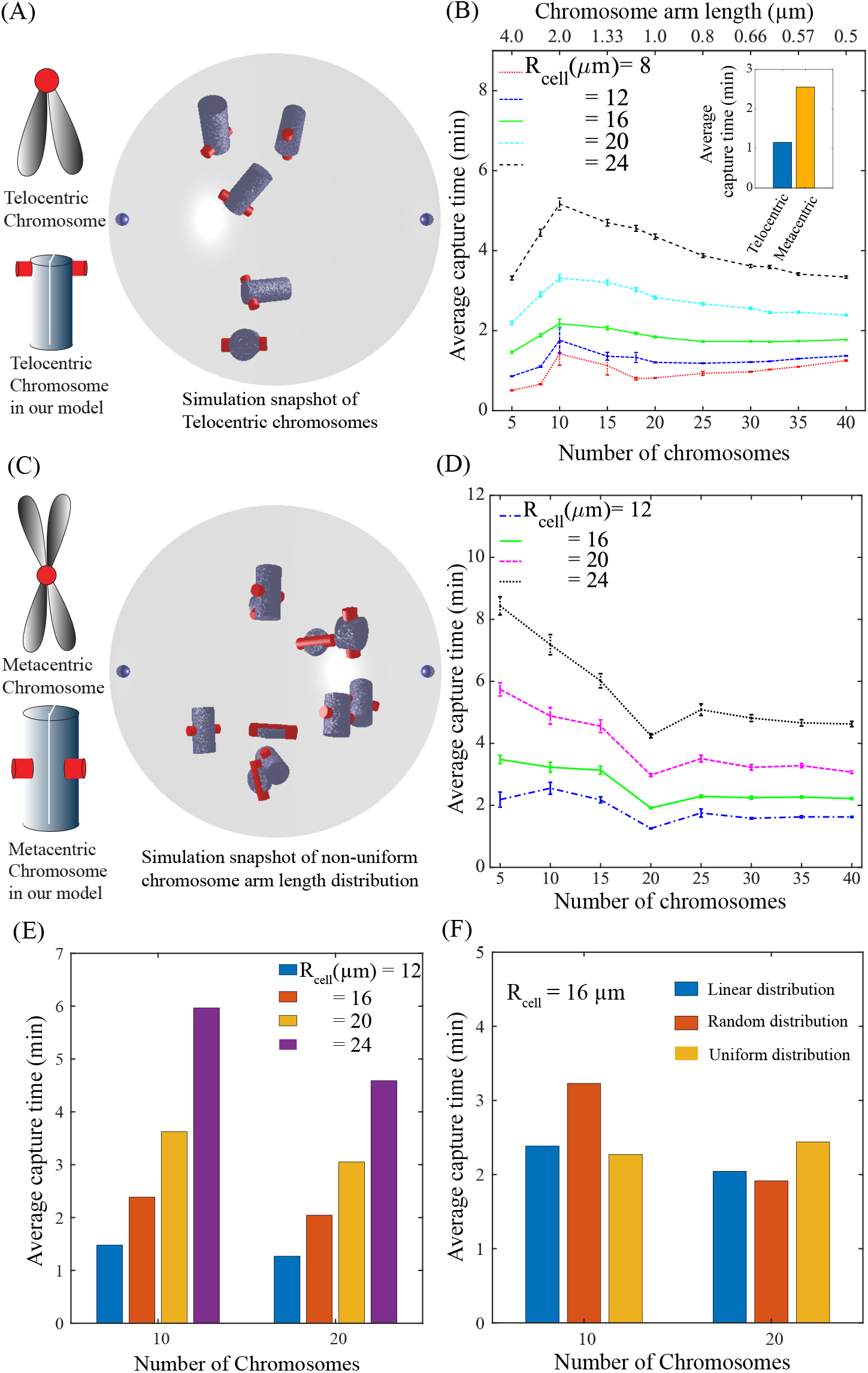
**(A)** Schematic of telocentric chromosome and simulation snapshot of a cell containing telocentric chromosomes.**(B)** Variation of *total capture time* with *chromosome number* for telocentric chromosomes with total chromosomes arm length conserved at 20 *μ*m. Chromosomes experience viscous drag inversely proportional to arm length. **Inset** contains capture time data for 5 chromosomes in a cell with radius 12 *μ*m performing only rotational diffusion (translational diffusion coefficient set to zero for both telocentric and metacentric chromosomes). **(C)** Schematic of metacentric chromosome and simulation snapshot of random distribution of chromosomal volume among 10 chromosomes. **(D)** Variation of capture time with chromosome numbers for randomly distributed (Gaussian distribution with mean 2 *μ*m and standard deviation 1.0 *μ*m.) chromosome arm lengths. The total chromosome arm length was conserved at 20 *μ*m. Chromosomes experience viscous drag inversely proportional to arm length. **(E)** Average capture time for chromosome numbers 10 and 20 where the individual chromosome arm lengths were increased linearly keeping the total chromosomal volume conserved. The chromosome arm lengths were increased linearly in the range 1.55 *μ*m to 2.45 *μ*m for 10 chromosomes and 0.55 *μ*m to 1.5 *μ*m for 20 chromosomes. **(F)** Comparison of capture times for linear distribution, random distribution (Gaussian distribution), and equal partitioning of the total chromosomal volume among the chromosomes. The distribution parameters are same as those applied in the simulations for Fig. 9D-9E.

### 6. Effect of random distribution of chromosome arm lengths

Chromosome sizes in a cell are seldom uniform. To check the effect of non-uniform distribution of the total chromosomal volume inside a cell, we simulated systems with a random distribution of the chromosome arm lengths keeping the total chromosomal volume conserved (see Fig. 9C). Here we considered a Gaussian distribution of chromosome arms with mean 2 *μ*m and standard deviation 1.0 *μ*m. The chromosomes experienced viscous drag inversely proportional to their arm lengths. Our results were qualitatively similar to those represented in Fig. 3D. When the chromosome numbers are small, many chromosomes are large and hence diffuse slowly. This leads to a comparatively larger capture time (Fig. 9D). As the number of chromosomes increased the chromosomes become smaller in size in order to maintain total chromo-somal volume conservation. This leads to a reduction in the viscous forces experienced by the chromosomes allowing them to diffuse faster. Accordingly, the capture time reduces with the increasing chromosome numbers similar to Fig. 3D. We also checked the capture times for 10 and 20 chromosomes when the chromosome arm length was increased linearly (Fig. 9E)). Note that the minimum arm length was 1.55 *μ*m for 10 chromosomes and 0.45 *μ*m for 20 chromosomes. This was motivated by the fact that the lengths of chromosomes of *S. Cerevisiae* show good linear fit (as given in the Saccharomyces Genome Database (SGD)) [68]. The capture time values were slightly lower than the values reported in Fig. 9D. This may be due to the fact that the random distribution of chromosomal volume leads to the presence of some large chromosomes which have higher shielding ability and low mobility leading to comparatively larger capture times. We compared the capture times for linear distribution, random distribution (Gaussian distribution), and equal partitioning of the total chromosomal volume among the chromosomes (Fig. 9F). Our data shows that linear distribution and equal partitioning of the chromosome volume lead to similar capture times. For fewer chromosomes, random distribution leads to larger capture times due to the presence of several large chromosomes.

## References

[1] Scholey JM, Brust-Mascher I, and Mogilner A. Cell division. Nature, 422:1–7, 2003.

[2] Alex Mogilner and Erin Craig. Towards a quantitative understanding of mitotic spindle assembly and mechanics. Journal of Cell Science, 123:3435–3445, 2010.

[3] R Heald and A Khodjakov. Thirty years of search and capture: The complex simplicity of mitotic spindle assembly. J. Cell Biol., 211(6):1103–1111, 2015.

[4] T E Holy and S Leibler. Dynamic instability of microtubules as an efficient way to search in space. PNAS, 91 (12):5682–5685, 1994.

[5] M Dogterom and S Leibler. Physical aspects of the growth and regulation of microtubule structures. Physical Review Letters, 70(9):1347–1350, 1993.

[6] T Mitchison and M Kirschner. Dynamic instability of microtubule growth. Nature, 312:237–242, 1984.

[7] M Kirschner and T Mitchison. Beyond self-assembly: From microtubules to morphogenesis. Cell, 45:329–342, 1986.

[8] T l Hill. Theoretical problems related to the attachment of microtubules to kinetochores. PNAS, 82:4404–4408, 1985.

[9] R. Wollman, E.N. Cytrynbaum, J.T. Jones, T. Meyer, J.M. Scholey, and A. Mogilner. Efficient chromosome capture requires a bias in the ‘search-and-capture’ process during mitotic-spindle assembly. Current Biology, 15:828–832, 2005.

[10] Raja Paul, Roy Wollman, William T. Silkworth, Isaac K. Nardi, Daniela Cimini, and Alex Mogilner. Computer simulations predict that chromosome movements and rotations accelerate mitotic spindle assembly without compromising accuracy. Proceedings of the National Academy of Sciences, 106(37):15708–15713, 2009.

[11] D Cimini and F Degrassi. Aneuploidy: a matter of bad connections. Trends Cell Biol., 15:442–451, 2005.

[12] CE Walczak, S Cai, and A Khodjakov. Mechanisms of chromosome behaviour during mitosis. Nat Rev Mol Cell Biol., 11:91–102, 2010.

[13] S Som, S Chatterjee, and R Paul. Mechanistic three-dimensional model to study centrosome positioning in the interphase cell. Phys. Rev. E, 99:012409, 2019.

[14] S. Chatterjee, A. Sarkar, J. Zhu, A. Khodjakov, Mogilner, and R. Paul. Mechanics of multicentrosomal clustering in bipolar mitotic spindles. Biophysical Journal, 119(2):434–447, 2020.

[15] Saptarshi Chatterjee, Subhendu Som, Neha Varshney, PVS Satyadev, Kaustuv Sanyal, and Raja Paul. Mechanics of microtubule organizing center clustering and spindle positioning in budding yeast cryptococcus neoformans. Phys. Rev. E, 104:034402, 2021.

[16] N Varshney, S Som, S Chatterjee, S Sridhar, D Bhattacharyya, R Paul, and K Sanyal. Spatio-temporal regulation of nuclear division by aurora b kinase ipl1 in cryptococcus neoformans. PLoS Genet, 15(2):e1007959, 2019.

[17] N. Pavin and I. M. Tolic. Self-organization and forces in the mitotic spindle. Annu Rev Biophys, 45:279–98, 2016.

[18] Manuel Théry, Andrea Jiménez-Dalmaroni, Victor Racine, Michel Bornens, and Frank Julicher. Experimental and theoretical study of mitotic spindle orientation. Nature, 447(7143):493–6, 2007.

[19] N. Minc, D. Burgess, and F. Chang. Influence of cell geometry on division-plane positioning. Cell, 144(3):414–26, 2011.

[20] G. Gay, T. Courtheoux, C. Reyes, S. Tournier, and Y. Gachet. A stochastic model of kinetochore-microtubule attachment accurately describes fission yeast chromosome segregation. J Cell Biol, 196(6):757–74, 2012.

[21] A. V. Zaytsev, D. Segura-Pena, M. Godzi, A. Calderon, E. R. Ballister, R. Stamatov, A. M. Mayo, L. Peterson, B. E. Black, F. I. Ataullakhanov, M. A. Lampson, and E. L. Grishchuk. Bistability of a coupled aurora b kinase-phosphatase system in cell division. Elife, 5:e10644, 2016.

[22] M. W. Elting, M. Prakash, D. B. Udy, and S. Dumont. Mapping load-bearing in the mammalian spindle reveals local kinetochore fiber anchorage that provides mechanical isolation and redundancy. Curr Biol, 27(14):2112– 2122.e5, 2017.

[23] J. Li, L. Cheng, and H. Jiang. Cell shape and intercellular adhesion regulate mitotic spindle orientation. Mol Biol Cell, 30(19):2458–2468, 2019.

[24] Gaelle Letort, Isma Bennabia, Serge Dmitrieffb, François Nedelec, Marie-Hélène Verlhaca, and Marie-Emilie Terreta. A computational model of the early stages of acentriolar meiotic spindle assembly. Mol Biol Cell., 30(7):863–875, 2019.

[25] A. R. Lamson, C. J. Edelmaier, M. A. Glaser, and M. D. Betterton. Theory of cytoskeletal reorganization during cross-linker-mediated mitotic spindle assembly. Biophys J, 116(9):1719–1731, 2019.

[26] Fioranna Renda, Valentin Magidson, Irina Tikhonenko, Rebecca Fisher, Christopher Miles, Alex Mogilner, and Alexey Khodjakov. Effects of malleable kinetochore morphology on measurements of intrakinetochore tension. Open Biol., 10:200101, 2020.

[27] Fioranna Renda, Christopher Miles, Irina Tikhonenko, Rebecca Fisher, Lina Carlini, Tarun M. Kapoor, Alex Mogilner, and Alexey Khodjakov. Non-centrosomal microtubules at kinetochores promote rapid chromosome biorientation during mitosis in human cells. Current Biology, 32 (5):1049–1063.e4, 2022.

[28] Blackwell R, Edelmaier C, Sweezy-Schindler O, Lamson A, Gergely ZR, and O’Toole E et al. Physical determinants of bipolar mitotic spindle assembly and stability in fission yeast. Sci Adv., 3(1):e1601603, 2017.

[29] Edelmaier CJ, Lamson AR, Gergely ZR, Ansari S, Blackwell R, and Richard McIntosh JR et al. Mechanisms of chromosome biorientation and bipolar spindle assembly analyzed by computational modeling. Elife, 116(9):1–48, 2020.

[30] Zaytsev A V. and Grishchuk EL. Basic mechanism for biorientation of mitotic chromosomes is provided by the kinetochore geometry and indiscriminate turnover of kinetochore microtubules. Mol Biol Cell., 26:3985–3998, 2015.

[31] V Magidson, R Paul, N Yang, J G. Ault, C B. O’Connell, I Tikhonenko, B F. McEwen, A Mogilner, and A Khodjakov. Adaptive changes in the kinetochore architecture facilitate proper spindle assembly. Nature Cell Biology, 17:1134–1144, 2015.

[32] N P. Ferenz, R Paul, C Fagerstrom, A Mogilner, and P Wadsworth. Dynein antagonizes eg5 by crosslinking and sliding antiparallel microtubules. Curr Biol., 19(21):1833–1838, 2009.

[33] V Magidson, C B. O’Connell, J Lončarek, R Paul, A Mogilner, and A Khodjakov. The spatial arrangement of chromosomes during prometaphase facilitates spindle assembly. Cell, 146:555–567, 2011.

[34] Indrani Nayak, Dibyendu Das, and Amitabha Nandi. Kinetochore capture by spindle microtubules: why fission yeast may prefer pivoting to search-and-capture. bioRxiv, page http://dx.doi.org/10.1101/673723, 2019.

[35] Iana Kalinina, Amitabha Nandi, Petrina Delivani, Mariola R. Chacón, Anna H. Klemm, Damien Ramunno-Johnson, Alexander Krull, Benjamin Lindner, Nenad Pavin, and Iva M. Toli ć -Nørrelykke. Pivoting of microtubules around the spindle pole accelerates kinetochore capture. Nat Cell Biol., 15(1):82–87, 2013.

[36] Kliuchnikov E, Zhmurov A, Marx KA, Mogilner A, and Barsegov V. Celldynamo–stochastic reaction-diffusion-dynamics model: Application to search-and-capture process of mitotic spindle assembly. PLoS Comput Biol, 18(6):e1010165, 2022.

[37] D.H. Wurster and K. Benirschke. Indian muntjac, muntiacus muntjak: a deer with a low diploid chromosome number. Science, 168:1364–1366, 1970.

[38] Vicky Tsipouri, Mary G. Schueler, Sufen Hu, Amalia Dutra, Evgenia Pak, Harold Riethman, and Eric D. Green. Comparative sequence analyses reveal sites of ancestral chromosomal fusions in the indian muntjac genome. Genome Biology, 9, 10 2008.

[39] Danica Drpic, Ana C. Almeida, Paulo Aguiar, Fioranna Renda, Joana Damas, Harris A. Lewin, Denis M. Larkin, Alexey Khodjakov, and Helder Maiato. Chromosome segregation is biased by kinetochore size. Current Biology, 28:1344–1356, 2018.

[40] Ana C. Almeida, Joana Soares de Oliveira, Danica Drpic, Liam P. Cheeseman, Joana Damas, Harris A. Lewin, Denis M. Larkin, Paulo Aguiar, António J. Pereira, and Helder Maiato. Augmin-dependent microtubule selforganization drives kinetochore fiber maturation in mammals. Cell articles, 39:110610, 2022.

[41] F Yang, P C M O’brien, J Wienberg, and M A Ferguson-Smith. A reappraisal of the tandem fusion theory of karyotype evolution in the indian muntjac using chromosome painting. Chromosome Research, 5:109–117, 1997.

[42] Jingchuan Luo, Xiaoji Sun, Brendan P. Cormack, and Jef D. Boeke. Karyotype engineering by chromosome fusion leads to reproductive isolation in yeast. Nature, 560:392–396, 8 2018.

[43] Yangyang Shao, Ning Lu, Zhenfang Wu, Chen Cai, Shanshan Wang, Ling Li Zhang, Fan Zhou, Shijun Xiao, Lin Liu, Xiaofei Zeng, Huajun Zheng, Chen Yang, Zhihu Zhao, Guoping Zhao, Jin Qiu Zhou, Xiaoli Xue, and Zhongjun Qin. Creating a functional single-chromosome yeast. Nature, 560:331–335, 8 2018.

[44] Kaihang Wang, Daniel De La Torre, Wesley E Robertson, and Jason W Chin. Programmed chromosome fission and fusion enable precise large-scale genome rearrangement and assembly downloaded from. Science, 2019.

[45] L. Fishman, J. H. Willis, C. A. Wu, and Y. W. Lee. Comparative linkage maps suggest that fission, not polyploidy, underlies near-doubling of chromosome number within monkeyflowers (mimulus; phrymaceae). Heredity, 112:562–568, 2014.

[46] Christine J Harrison, Terence D Allen’, Martin Britch, and Rodney Harris. High-resolution scanning electron microscopy of human metaphase chromosomes. J. Cell Sci, 56:409–422, 1982.

[47] A t Sumner. Scanning electron microscopy of mammalian chromosomes from prophase to telophase. Chromosoma, 100:410–418, 1991.

[48] Helder Maiato, Conly L. Rieder, and Alexey Khodjakov. Kinetochore-driven formation of kinetochore fibers contributes to spindle assembly during animal mitosis. Journal of Cell Biology, 167:831–840, 12 2004.

[49] U. Serdar Tulu, Carey Fagerstrom, Nick P. Ferenz, and Patricia Wadsworth. Molecular requirements for kinetochore-associated microtubule formation in mammalian cells. Current Biology, 16:536–541, 3 2006.

[50] Evgenii Kliuchnikov, Kenneth A Marx, Alex Mogilner, and Valeri Barsegov. Interrelated effects of chromosome size, mechanics, number, location-orientation and polar ejection force on the spindle accuracy: a 3d computational study. Mol Biol Cell., 2023.

[51] Helder Maiato and Sónia Silva. Double-checking chromosome segregation. J Cell Biol, 222 (5):e202301106, 2023.

[52] Kenneth H. Wolfe. Origin of the yeast whole-genome duplication. PLoS Biology, 13, 2015.

[53] Yuuta Moriyama and Kazuko Koshiba-Takeuchi. Significance of whole-genome duplications on the emergence of evolutionary novelties. Briefings in Functional Genomics, 17:329–338, 9 2018.

[54] Kenneth H. Wolfe and Denis C. Shields. Molecular evidence for an ancient duplication of the entire yeast genome. Nature, 387:708–713, 1997.

[55] Francisco J Ayala and Mario Coluzzi. Chromosome speciation: Humans, drosophila, and mosquitoes. PNAS, 2005.

[56] Raúl A. Ortiz-Merino, Nurzhan Kuanyshev, Stephanie Braun-Galleani, Kevin P. Byrne, Danilo Porro, Paola Branduardi, and Kenneth H. Wolfe. Evolutionary restoration of fertility in an interspecies hybrid yeast, by whole-genome duplication after a failed mating-type switch. PLoS Biology, 15, 5 2017.

[57] Manolis Kellis, Bruce W Birren, and Eric S Lander. Proof and evolutionary analysis of ancient genome duplication in the yeast saccharomyces cerevisiae. Nature, 2004.

[58] Camille Berthelot, F. édéric Brunet, Domitille Chalopin, A. élie Juanchich, Maria Bernard, Benjamin Noël, Pascal Bento, Corinne Da Silva, Karine Labadie, Adriana Alberti, Jean Marc Aury, Alexandra Louis, Patrice Dehais, Philippe Bardou, Jérôme Montfort, Christophe Klopp, Cédric Cabau, Christine Gaspin, Gary H. Thorgaard, Mekki Boussaha, Edwige Quillet, René Guyomard, Delphine Galiana, Julien Bobe, Jean Nicolas Volff, Carine Genet, Patrick Wincker, Olivier Jaillon, Hugues Roest Crollius, and Yann Guiguen. The rainbow trout genome provides novel insights into evolution after whole-genome duplication in vertebrates. Nature Communications, 5, 4 2014.

[59] Jurriaan M. De Vos, Hannah Augustijnen, Livio Bätscher, and Kay Lucek. Speciation through chromosomal fusion and fission in lepidoptera: Chromosomal fusion & fission. Philosophical Transactions of the Royal Society B: Biological Sciences, 375, 8 2020.

[60] Ryan J. Quinton, Amanda DiDomizio, Marc A. Vittoria, Kristýna Kotýnková, Carlos J. Ticas, Sheena Patel, Yusuke Koga, Jasmine Vakhshoorzadeh, Nicole Hermance, Taruho S. Kuroda, Neha Parulekar, Alison M. Taylor, Amity L. Manning, Joshua D. Campbell, and Neil J. Ganem. Whole-genome doubling confers unique genetic vulnerabilities on tumour cells. Nature, 590:492–497, 2 2021.

[61] Rachel Newcomb, Emily Dean, Brock J. McKinney, and James V. Alvarez. Context-dependent effects of wholegenome duplication during mammary tumor recurrence. Scientific articles, 11, 12 2021.

[62] Saioa López et. al. Interplay between whole-genome doubling and the accumulation of deleterious alterations in cancer evolution. Nature Genetics, 52:283–293, 3 2020.

[63] Jonathan L. Gordon, Kevin P. Byrne, and Kenneth H. Wolfe. Mechanisms of chromosome number evolution in yeast. PLoS Genetics, 7, 7 2011.

[64] Ronald D Vale. Review the molecular motor toolbox for intracellular transport transport involves molecular motor proteins that carry cargo directionally along a cytoskeletal track (myosins along actin and kinesins and dyneins along microtu. Cell, 112:467–480, 2003.

[65] D J Palmer, J B Helms, C J Becker, L Orci, J E Rothman, Richard B Vallee, and Michael P Sheetz. 47. c. d’souza-schorey. Bio-chem. Biophys. Res. Commun, 6:14, 1994.

[66] Jacob A. Herman, Matthew P. Miller, and Sue Biggins. Chtog is a conserved mitotic error correction factor. eLife, 9:1–28, 12 2020.

[67] Michael A. Lampson and Ekaterina L. Grishchuk. Mechanisms to avoid and correct erroneous kinetochore-microtubule attachments. Biology, 6, 3 2017.

[68] Reference genome: Saccharomyces cerevisiae s288c. Saccharomyces Genome Database, https://www.yeastgenome.org/.

